# Spike Detection Using FRI Methods and Protein Calcium Sensors: Performance Analysis and Comparisons

**DOI:** 10.1101/029124

**Authors:** Stephanie Reynolds, Jon Oñativia, Caroline S Copeland, Simon R Schultz, Pier Luigi Dragotti

## Abstract

Fast and accurate detection of action potentials from neurophysiological data is key to the study of information processing in the nervous system. Previous work has shown that finite rate of innovation (FRI) theory can be used to successfully reconstruct spike trains from noisy calcium imaging data. This is due to the fact that calcium imaging data can be modeled as streams of decaying exponentials which are a subclass of FRI signals. Recent progress in the development of genetically encoded calcium indicators (GECIs) has produced protein calcium sensors that exceed the sensitivity of the synthetic dyes traditionally used in calcium imaging experiments. In this paper, we compare the suitability for spike detection of the kinetics of a new family of GECIs (the GCaMP6 family) with the synthetic dye Oregon Green BAPTA-1. We demonstrate the high performance of the FRI algorithm on surrogate data for each calcium indicator and we calculate the Cramér-Rao lower bound on the uncertainty of the position of a detected spike in calcium imaging data for each calcium indicator.

## I. Introduction

The firing of action potentials transmits information in neuronal networks. In order to study information processing in the nervous system, it is important to be able to detect accurately the time points of action potentials (spikes) across populations of neurons from neurophysiological data. As the concentration of intracellular free calcium is a reliable indicator of neuronal activity in many cell types, several optical imaging methods rely on fluorescent indicators which report calcium concentration changes via changes in their intensity (see [1] for a review).

Throughout the past three decades synthetic dyes have been predominantly used as the fluorescent indicator in calcium imaging experiments, with Oregon Green BAPTA-1 (OGB-1) and fluo-4 among the most commonly used [2]. However, a new family of protein calcium sensors (GCaMP6) has recently been engineered by Chen et al. [3] and has been found to exceed the sensitivity of synthetic dyes, an achievement which no other protein calcium sensor has yet accomplished. Thestrup et al.’s ‘Twitch’ sensors, a family of optimised ratiometric calcium indicators, further represent the recent progress in the engineering of protein calcium sensors [4]. Unlike synthetic dyes, protein calcium sensors are able to selectively label cell populations and can be used for chronic *in vivo* imaging. These advantages may result in protein calcium sensors such as GCaMP6 and Twitch being preferred as fluorescent indicators in future calcium imaging work.

Several approaches have been taken to detect spikes in calcium imaging data. Sasaki et al. developed a supervised machine learning algorithm that utilises principal component analysis on calcium imaging data to detect spikes [5]. Similarly, in [6] Vogelstein et al. introduce an algorithm that learns parameters from calcium imaging data and then performs an approximate *maximum a posteriori* estimation to infer the most likely spike train given fluorescence data. In [7] Grewe et al. employ an algorithm which detects events via a combination of amplitude thresholding and analysis of the fluorescence sequence integral. In [8], Schultz et al. use an event template derived from imaging data to identify the spike train which correlates most highly with the noisy fluorescence signal.

In this study we seek to compare the suitability for spike detection from calcium imaging data of the kinetics of the synthetic dye OGB-1 and two GCaMP6 sensors (the fast variant GCaMP6f and the slow variant GCaMP6s). The pulse in a neuron’s localised fluorescence data that occurs as a result of the firing of an action potential has a different characteristic shape for each of the fluorescent indicators we consider (GCaMP6f, GCaMP6s and OGB-1). By incorporating these characteristic pulse shapes into our model of imaging data for each fluorescent indicator, and simulating surrogate data based on these models, we wish to compare spike detection performance on each surrogate data set. Moreover, we calculate the Cramér-Rao bound for the uncertainty of the estimated location of a spike in calcium imaging data for each fluorescent indicator and compare the spike detection algorithm’s performance against this lower bound.

To make our comparisons of the indicators we use the finite rate of innovation (FRI) spike detection algorithm developed by Oñativia et al. in [9]. This algorithm exploits the fact that calcium imaging data can be modelled as streams of decaying exponentials, which are a subclass of FRI signals [10]. This allows the authors to apply FRI methods (for an overview see [11]) to reconstruct the signal from noisy data. The algorithm, which can be performed in real-time, was shown to have a high spike detection rate and low false positive rate on both real and surrogate data.

This paper is organised as follows: in Section II we formulate mathematically the problem of spike detection from calcium imaging data. In Section III we provide a review of Oñativia et al.’s FRI spike detection algorithm and present our method of generating surrogate calcium imaging data. In Section IV we formulate an expression for the Cramér-Rao bound on the uncertainty of the position of one spike in calcium imaging data. We then show results from our simulations and compare Cramér-Rao bounds for imaging data from different fluorescent indicators in Section V. Finally, in Section VI we conclude.

## II. Problem Formulation

Each action potential produces a characteristic pulse shape in the corresponding neuron’s fluorescence signal. We therefore model the fluorescence signal of one neuron over time as a convolution of that neuron’s spike train and the characteristic pulse shape, such that

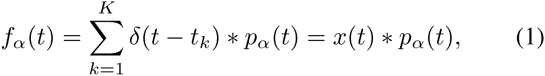

where 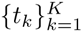 are the time points of the *K* spikes and the spike train is written 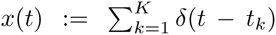. The characteristic pulse shape of the fluorescence signal differs for each fluorescent indicator due to the differences in the indicators’ kinetics. We make the assumptions that each of the three pulse shapes have an instantaneous rise and exponential decay, where the speed of the decay (*α*) and peak amplitude (*A*) are different for each indicator, such that

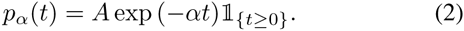

## III. Methods

### A. Finite rate of innovation applied to spike detection

In [9], Oñativia et al. develop an FRI algorithm for spike detection in two-photon calcium imaging data. A review of that algorithm, which we use to detect spikes in our surrogate data, is provided here. We refer readers to Oñativia et al. for further detail. We start with a useful definition.

#### Definition III.1.

An exponential reproducing kernel is one such that, when summated with its shifted versions, it generates exponentials of the form exp (*γ_m_t*):

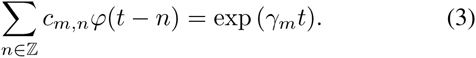

The values *c_m,n_* are referred to as the coefficients of the exponential reproducing kernel. As our sampling kernel we use an E-spline, which is a type of exponential reproducing kernel that has compact support (see [12], [13] for more details).

Initially, the fluorescence signal *f_α_*(*t*) is filtered with an exponential reproducing kernel 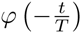 and sampled *N* times with sampling period *T*, such that

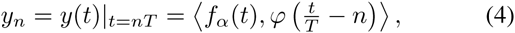

for *n* ∈ {0, 1, &, *N* −1}. Finite differences are then computed to form samples

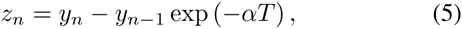

for *n* ∈{1, 2, &, *N* − 1}. Using Parseval’s theorem, it is possible to show that the computation of *z_n_* in Equation (5) is equivalent to filtering the spike train 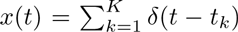 with a new exponential reproducing kernel 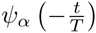, such that

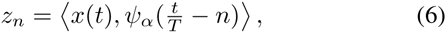

where *ψ_α_*(*t*)= *β_αT_* (−*t*) ∗ *ϕ*(*t*) and *β_αT_* (*t*) is a first-order E-spline (for further details see [12]). The initial exponential reproducing kernel *ϕ* is chosen to ensure that *ψ_α_*(*t*) reproduces exponentials with exponents of the form

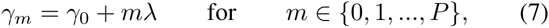

where, in this case, we choose 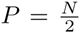. Ensuring exponents of this form allows us to apply the annihilating filter method to the exponential sample moments

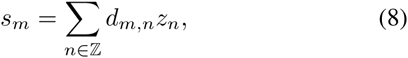

where *d_m,n_* are the coefficients of the exponential reproducing kernel *ψ_α_*(*t*). With exponents of the form in (7), we can rearrange Equation (8) to become

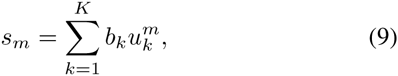

where 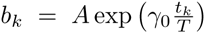 and 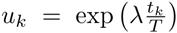. We then construct a Toeplitz matrix S from the exponential sample moments *s_m_*. In the idealised, noiseless scenario in which we know the value of *K*, we can use de Prony’s method along with the condition that *h*_0_ = 1 to find the unique annihilating filter h such that

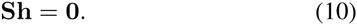

Given the value of h, we can calculate the zeroes of its z-transform, which are equivalent to 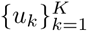. From these values we can retrieve the spike times 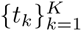.

In real data, we do not know the value of *K* and must estimate it from noisy samples. Oñativia et al.’s algorithm uses a double consistency approach to do this. Firstly, for each consecutive 32-sample window contained within the data, the value of *K* in that window is estimated from the singular value decomposition of S. The corresponding spikes are then estimated using the annihilating filter method outlined above.

Secondly, it is assumed that there is a single spike in each consecutive 8-sample window contained within the data, and the position of that spike is estimated using the annihilating filter method. A joint histogram is constructed, containing all the estimated spikes and their position within the data. The peaks of this histogram, corresponding to spikes which were consistently estimated across windows, are selected as the positions of the true spikes.

### B. Generating surrogate data

We assume that the occurrence of spikes follows a Poisson distribution with parameter *λ* (spikes per second). We use the spike rate parameter *λ* =0.25Hz, which corresponds to the experimentally measured spontaneous spike rate in the barrel cortex [14], as our further work will involve calcium imaging data from this brain region. To generate the spike train we use the fact that the waiting time between Poisson(*λ*) occurrences follows an exponential distribution with parameter *λ*.

For each fluorescent indicator, using the same generated spike times 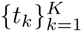, we calculate a fluorescence waveform

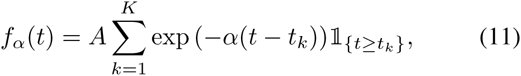

where the parameters *α* and *A* are specific to the fluorescent indicator used. The values for each *α* and *A* were derived from experimental results given in [3] and can be found in Table I. We sample each fluorescence waveform *N* times at time resolution *T* and add white noise, such that we have samples

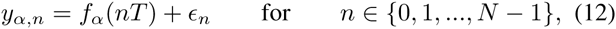

with 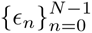 a family of independent and identically distributed Gaussian random variables with variance *σ*^2^. Using this method we have, for each fluorescent indicator, a set of noisy samples of the fluorescence signal. The performance of the algorithm on surrogate data for each fluorescent indicator is directly comparable as the same spike train and noise realisations have been used.

**TABLE I.** Parameters for Calcium Transient Model.

| Calcium indicator | A | $\alpha$ |
| --- | --- | --- |
| GCaMP6f | 0.19 | $\frac{\log(2)}{0.142}$ |
| GCaMP6s | 0.23 | $\frac{\log(2)}{0.55}$ |
| OGB-1 | 0.1642 | $\frac{1}{0.581}$ |

## IV. Derivation of the CramÉr-Rao Bound

We calculate the Cramér-Rao bound for the uncertainty of the estimated location of a spike in a noisy fluorescence signal. We consider the fluorescence signal from Equation (1) in the case that *K* =1, so that *f_α_*(*t*) becomes

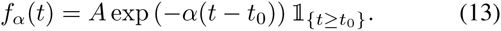

The signal *f_α_*(*t*) is then filtered with function 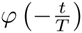 and sampled *N* times with time resolution *T*, such that we obtain the noisy samples

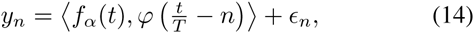

for *n* ∈ {0, 1, &, *N* − 1}, where 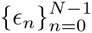 are independent and identically distributed Gaussian random variables with variance *σ*^2^.

### Proposition IV.1.

*The Cramér-Rao bound of estimating* 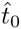 *from N noisy samples of the form* 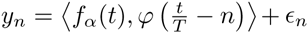*, where* 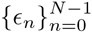 *is a family of independently and identically Gaussian distributed noise, is*

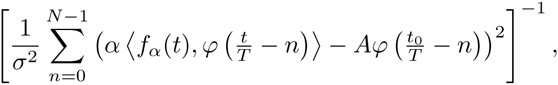

*where σ*^2^ *is the variance of ε_n_ for n* ∈ {0, 1, &, *N* − 1}.

#### Proof

We write 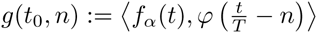, so that

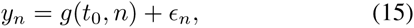

for *n* ∈{0, 1, &, *N* −1}. The Fisher Information for estimating a single parameter 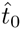 from *N* samples {*y*_0_*, y*_1_, &, *y_N_*_−1_} with i.i.d. Gaussian noise is

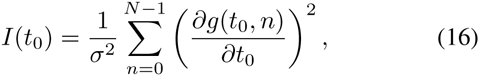

where *σ*^2^ is the variance of the noise. We have that

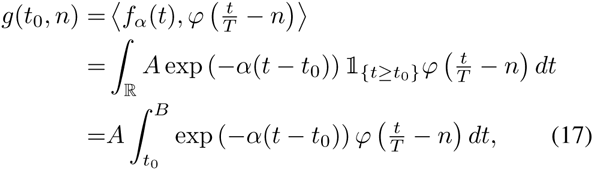

where B is the upper limit of the support of the kernel *ϕ*, we know our kernel has finite support as it is an E-spline. Denoting the integrand as *h*(*t, t*_0_), we have

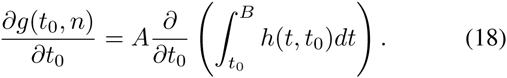

By the Leibniz Integral Rule, as 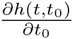 exists and is continuous in *t*_0_, we can write

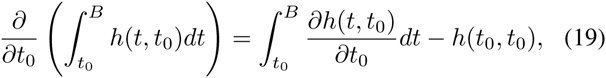

from which it follows that

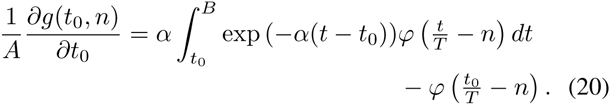

This reduces to

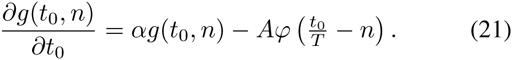

Plugging Equation (21) into Equation (16), we arrive at the statement of the proposition.

## V. Results

### A. Performance analysis based on theoretical bounds

For each fluorescent indicator, we calculated the Cramér-Rao lower bound (CRB) for the uncertainty of estimating the position of one spike from noisy samples of fluorescence sequence data. We compared this to the mean square error between the position of the real spike and the position estimated by the FRI spike detection algorithm. In particular, we assumed there was one spike in a time interval of length 1 second and that samples were obtained in the manner described in Section III with sampling rate 16Hz. The performance of the algorithm compared to the CRB was computed for a range of values of the variance of the additive Gaussian noise (*σ*^2^, see Equation (12)). For each value of *σ*^2^, the results were averaged over 1000 realisations of noise.

As can be seen in Figure 1a, for each characteristic pulse shape, the FRI algorithm achieves near optimal performance for noise variances beneath a certain break point. This break point is reached first for spikes with a GCaMP6s characteristic pulse shape.

**Fig. 1.**
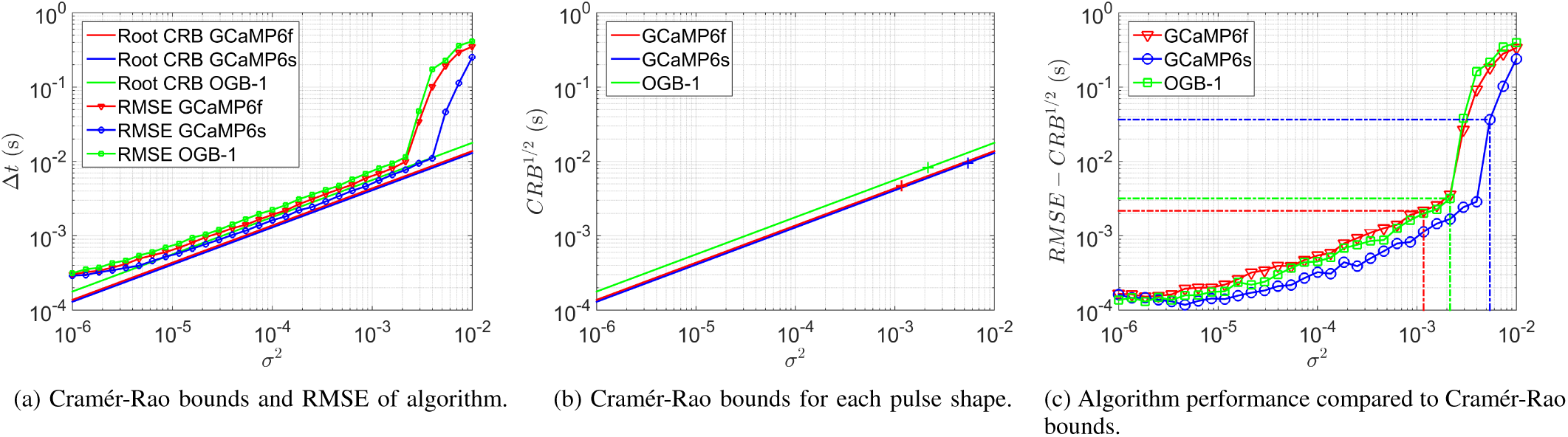
FRI algorithm performance compared to Cramér-Rao bounds for surrogate data with 16Hz sampling rate. Results are averaged over 1000 realisations of noise. (a) The variance of the additive Gaussian noise (*σ*^2^) is plotted against the root mean square error (RMSE) of the FRI algorithm and the square root of the Cramer-Rao bound for each characteristic pulse shape. (b) The noise variance *σ*^2^ is plotted against the root of the Cramér-Rao bound. The crosses indicate the point on each indicator’s curve which corresponds to an SNR of 10dB. (c) The residue of the RMSE from the root of the Cramér-Rao bound is plotted against the noise variance *σ*^2^. The dashed lines identify the point corresponding to an SNR of 10dB for each fluorescent indicator.

GCaMP6s spikes have the lowest theoretical lower-bound on uncertainty for all values of *σ*^2^, whereas OGB-1 spikes have the highest, this is illustrated in Figure 1b. Furthermore, as shown in Figure 1c, the difference between the root mean square error (RMSE) of the spike detection algorithm and the theoretical lower bound of performance (the square root of the CRB) is significantly smaller when detecting a GCaMP6s characteristic pulse shape than for GCaMP6f and OGB-1 pulse shapes.

The above analysis indicates that, under the same imaging conditions (and therefore the same noise variance), the FRI algorithm locates a spike with the least uncertainty when that spike has a GCaMP6s pulse shape. This can be attributed to the fact that GCaMP6s spikes have the highest operating signal-to-noise ratio (SNR), a property which arises from their relatively high peak amplitude (see Table I). When we compare algorithm performance against SNR instead of noise variance, we effectively normalise the amplitudes of the pulses. It can be seen in Figure 2 that, for each SNR, the CRB and RMSE are lowest for a GCaMP6f characteristic pulse shape.

**Fig. 2.**
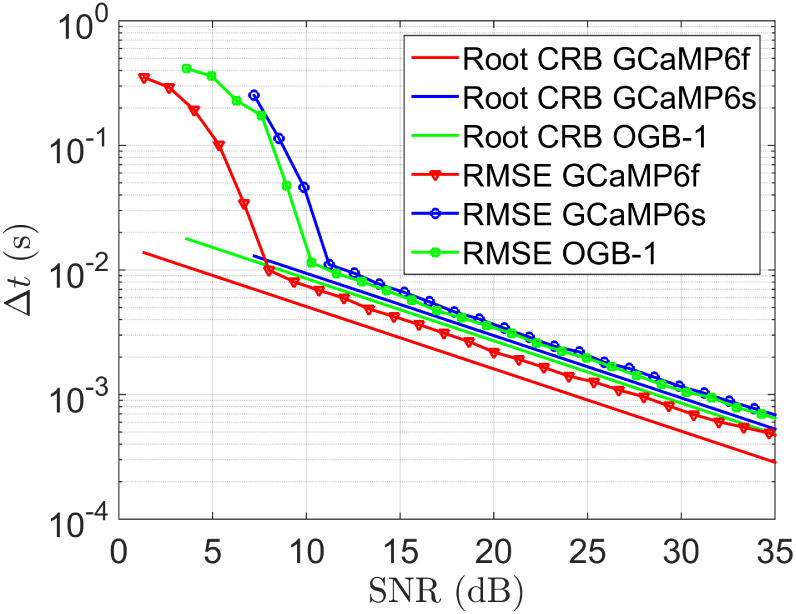
FRI algorithm performance compared to Cramér-Rao bounds for surrogate data with 16Hz sampling rate. Results are averaged over 1000 realisations of noise traces. Signal-to-noise ratio is plotted against the root mean square error (RMSE) of the FRI algorithm and the square root of the Cramér-Rao bound for each characteristic pulse shape.

Our performance analysis showed that, under the same imaging conditions, the lowest uncertainty on the position of the detected spike is achieved when that spike has a GCaMP6s characteristic pulse shape. The high relative performance of GCaMP6s is likely to stem from its advantage of having the highest peak amplitude, and thus the highest operating signal-to-noise ratio. When the uncertainty in the position of the detected spike is compared across the same signal-to-noise ratios, therefore removing the impact of the discrepancy in peak amplitudes, the GCaMP6 variant with the fastest pulse decay (GCaMP6f) performs best.

### B. Simulation results

In the manner described in Section III-B, we simulated surrogate fluorescence sequence data for 20 pairings of noise variance (*σ*^2^) and sampling rate. The algorithm performance on the GCaMP6f, GCaMP6s and OGB-1 surrogate data is directly comparable as the same simulated noise traces and spike trains were used. The following performance statistics are averaged over 100 realisations for each *σ*^2^ and sampling rate pairing.

The FRI algorithm achieved a high spike detection rate on surrogate data for each fluorescent indicator, regularly detecting above 90% of spikes. Table II shows the mean and standard deviation of the spike detection rate across surrogate data for each fluorescent indicator. From these data it can be seen that, for noise variances beneath 8 10^−4^, there was little difference in the ability of the FRI algorithm to detect spikes with the three different pulse shapes.

**TABLE II.** Average spike detection rate (%) of FRI algorithm on surrogate data (mean ± standard deviation across indicators)

| $\sigma^2$ | Sampling rate (Hz) | | | | |
| --- | --- | --- | --- | --- | --- |
|  | 8 | 16 | 32 | 64 | 128 |
| $8 \cdot 10^{-4}$ | 85.9±6.8 | 91.7±4.7 | 93.2±3.0 | 91.3±5.7 | 90.3±4.7 |
| $3 \cdot 10^{-4}$ | 88.6±2.2 | 93.7±1.0 | 95.1±1.6 | 94.4±2.1 | 92.8±2.7 |
| $3 \cdot 10^{-5}$ | 89.6±0.5 | 94.3±0.6 | 96.5±0.2 | 96.2±0.9 | 96.0±1.7 |
| $3 \cdot 10^{-6}$ | 90.7±0.9 | 94.4±0.4 | 96.0±0.6 | 95.6±1.3 | 96.2±1.4 |

No false positives were produced in 81% of surrogate data sequences. For a spike rate of 0.25Hz, the corresponding false positive rates were 8.7 10^−3^ Hz, 3.4 10^−4^ Hz and 7.1 10^−3^ Hz for pulses with OGB-1, GCaMP6f and GCaMP6s characteristic pulse shapes, respectively.

For a given power of noise, the three indicators have different operating SNRs. For example, the corresponding average SNR for a noise power of 8 10^−4^ is 5dB, 1dB and 9.5dB for OGB-1, GCaMP6f and GCaMP6s pulses, respectively. If the comparison of spike detection performance is made linearly in terms of SNR it can be seen that the fastest decaying pulse GCaMP6f outperforms both OGB-1 and GCaMP6s (by average spike detection rate margins of 2.1% and 4.3%, respectively).

The FRI algorithm produces high spike detection rates and low false positive rates on surrogate data for each of the three fluorescent indicators. The performance on each surrogate data set under the same imaging conditions (same *σ*^2^ and sampling rate) was very similar. When normalising the SNR and comparing spike detection performance, it was seen that the fast GCaMP6 variant outperformed the other two indicators, particularly at low SNRs.

## VI. Conclusion

We investigated the relative suitability for spike detection from calcium imaging data of three fluorescent indicators: the conventionally used synthetic dye OGB-1 and two new protein calcium sensors GCaMP6f and GCaMP6s. We demonstrated that the FRI algorithm achieves high spike detection rates and low false positive rates on surrogate data for each fluorescent indicator. Furthermore, we calculated an expression for the Cramér-Rao lower bound on the uncertainty of the estimated position of a spike in calcium imaging data and compared these bounds with the FRI algorithm’s performance on surrogate data. We found that, under the same imaging conditions, spikes with a GCaMP6s pulse shape have the lowest Cramér-Rao bound and the FRI algorithm is the closest to attaining that bound when detecting GCaMP6s spikes. The superiority, in this respect, of GCaMP6s can be attributed to the fact that, under the same imaging conditions as other indicators, the GCaMP6s pulse has a higher SNR. When assessing indicator performance linearly across SNRs, it is the indicator with the fastest decay (GCaMP6f) that performs the best.

